# High Content System for Quantifying Mitochondrial Morphology in Patient-Derived Human Cells

**DOI:** 10.1101/2025.05.08.652922

**Authors:** Kaitlyn Deschamps, Samantha Laville-Dupuy, Christina Yi Peng, Ray Truant

## Abstract

Mitochondrial function is critical for cellular health, with dysfunction contributing to human diseases. Structural changes in mitochondria, such as size and shape, reflect alterations in bioenergetics, fission-fusion dynamics, and metabolic homeostasis. Existing morphological quantification is outdated, can be biased, technologically limited, or overly complex. This study presents a high content system for quantifying morphology using open-access resources and widely available equipment. Fibroblasts were stained with PKmito^TM^ Dye Deep Red, imaged via automated confocal microscopy, and analyzed with CellProfiler™ and KNIME^®^. We tested different imaging conditions and found live-cell confocal imaging at 60x magnification provided the most precise measurements. Using this system, we found that human Huntington Disease fibroblast mitochondria were significantly smaller and more circular, suggesting increased fission. To confirm our results, we employed other mitochondrial assays and found elevated expression of the fission protein Drp1, reduced respiration, impaired iron uptake, and increased membrane potential. This system offers a robust, unbiased high content approach to studying mitochondrial morphology in disease.

## Introduction

Mitochondria are highly dynamic organelles that are critical for energy production, metabolic regulation, and iron homeostasis.^1, 2^ Mitochondria adapt to metabolic changes and energy demands by undergoing fission and fusion, ultimately governing ATP generation and health.^2–5^ Studying mitochondrial morphology and function has gained increasing attention due to their critical role in cellular physiology and disease mechanisms. However, much of the early experimental work was founded on now outdated, low-throughput methodologies and technologies with sampling designs of limited scope and rigour that may involve potential bias and investigator subjectivity.^6, 7^

One of the most widely used methods for assessing mitochondrial morphology is fluorescent staining with MitoTracker^TM^ Red. This carbocyanine-based dye passively diffuses into active mitochondria in a membrane potential-dependent manner.^8^ Despite its popularity, investigators using MitoTracker^TM^ products report one major limitation: phototoxicity caused by extensive illumination, which can adversely affect mitochondrial health and function.^9^

To overcome the toxicity of MitoTracker^TM^ Red, researchers from the Chen Lab at Peking University recently developed two alternative probes: PKmito^TM^ Dyes Red (PKMR) and Deep Red (PKMDR).^9^ These mitochondrial probes are similarly membrane potential-dependent, but are cyclooctatetraene-conjugated and explicitly developed for live-cell imaging. These dyes significantly reduce imaging-generated reactive oxygen species by utilizing triplet-state quenching, thereby minimizing phototoxicity.^9^ PKmito^TM^ Dyes offer improved stability and compatibility for long-term imaging studies, which is required for advancing research into mitochondrial morphology. While these imaging tools enhance our ability to quantify mitochondrial abnormalities, understanding dysfunction accurately in the context of disease requires clinically relevant human samples.

Studying mitochondrial dysfunction in neurodegenerative disorders like Huntington Disease (HD) has historically relied on animal models. Transgenic mouse models often express only a fragment of the disease-associated protein or contain mutation lengths that exceed what is commonly observed in patients.^10–12^ This disconnect raised concerns about their translational capacity to human pathology.^10, 11, 13–16^ To address these concerns, the Truant lab immortalized patient-derived fibroblasts using hTERT (TruHD cells; Table 1), which more accurately reflect human HD pathology.^17–20^

**Table 1.** Characterization of TruHD patient-derived fibroblasts.

| **Name and Primary ID (Coriell Institute)** | **CAG Length Allele 1** | **CAG Length Allele 2** | **Gender** | **CAP Score\*** | **Authentication** | **Mycoplasma Status** |
| --- | --- | --- | --- | --- | --- | --- |
| Control ND30014 | 21 | 18 | Female | N/A | PCR allele analysis | Negative |
| HD ND30013 | 43 | 17 | Male | 108 | PCR allele analysis | Negative |

\*CAG-Age product (CAP) Score = Age x (CAG – centering constant)/scaling constant. A score of 100 or higher indicates disease onset. Where less than 290=early progression; 290-367=mid progression; higher than 267=late progression.<sup>93</sup>

HD is caused by a CAG repeat expansion in the huntingtin gene, but additional factors such as DNA repair mechanisms and mitochondrial energetics, can influence the severity of the disease.^21, 22^ Mitochondrial dysfunction is widely accepted as a contributor to HD pathogenesis; however, the specific morphological and functional differences remain unclear.^7, 23–25^ It is widely accepted that mitochondria in HD patients demonstrate significant alterations in size and shape, but we anticipate more subtle differences.^6, 7^ HD is typically an adult-onset disease despite the presence of the gene mutation from birth, suggesting that dysfunction accumulates gradually over decades before reaching a pathogenic threshold. As a result, extreme abnormalities in mitochondrial morphology reported previously may be less distinct in this study. Using cells from HD patients with clinically relevant allele lengths provides a more accurate representation of disease and is invaluable for understanding the role of mitochondrial-related dysfunction in HD.

Here, we introduce a universal, high content method of assessing mitochondrial morphology using advanced live-cell imaging techniques. Our approach combines high-resolution imaging with quantitative analytical pipelines developed in CellProfiler^TM^ and KNIME^®^, allowing for accurate and replicable object identification, automated data management, and reproducible analysis. These pipelines can be adapted for any experimental techniques that quantify mitochondrial morphology. The accuracy of this live-cell mitochondrial analysis was assessed using orthogonal assays to measure respiration, Drp1 fission protein concentration, membrane potential, and transferrin levels in the context of TruHD fibroblasts. This study establishes a foundation for reliable and relevant investigations into mitochondrial dysfunction by refining the quantification of mitochondrial morphology.

## Results

A complete list of reagents and chemicals is listed in Table 2. Absolute values and standard deviations of all experimental conditions can be found in Table 3. Raw data are available through McMaster Dataverse, a collection within Borealis, the Canadian Dataverse Repository using https://doi.org/10.5683/SP3/VII0UC.26

**Table 2.** List of reagents, chemicals, and technology used.

| **Name** | **Source** | **Catalogue Code** | **Solubilization, Dilution, & Other** |
| --- | --- | --- | --- |
| LookOut<sup>®</sup> Mycoplasma PCR Detection Kit | Millipore Sigma | MP0035 |  |
| Phenol-red MEM | Gibco | 10370021 |  |
| Fetal bovine serum | Life Technologies | 098450 |  |
| GlutaMAX<sup>™</sup> | Thermo Fisher Scientific | 35050061 |  |
| CellCarrier-96 Ultra Microplates | Cell-Vis | P96-1.5H-N |  |
| PKmito Red and Deep Red* | Cytoskeleton Inc. | CY-SC052<br>CY-SC055 | 100% Dimethyl sulfoxide (DMSO) |
| NucBlue<sup>™</sup> Live Cell Stain ReadyProbes<sup>™</sup> reagent | Invitrogen | REF R37605 | 1 drop/mL |
| Phenol red-free MEM | Gibco<sup>™</sup> | 51200038 |  |
| Triton<sup>™</sup> X-100 | BioShop<sup>®</sup> Canada | TRX777.500 | 0.2% |
| Anti-DRP1 rabbit monoclonal antibody | Cell Signaling Technology | (D6C7) #8570 | 1:100 |
| Cy5<sup>®</sup> goat anti-rabbit IgG (H+L) | Life Technologies | A10523 | 1:500 |
| Hoechst 33258 | Life Technologies | H3569 | 0.2 µg/mL |
| 8-well XFp PDL Miniplate | Agilent Technologies | 103748-000 |  |
| XFp Extracellular Flux Cartridges | Agilent Technologies | 102983-000 |  |
| Seahorse XF DMEM Assay Medium pH 7.4 | Agilent Technologies | 103575-100 |  |
| Glucose | Agilent Technologies | 103577-100 | 10 mM |
| Glutamine | Agilent Technologies | 103579-100 | 2 mM |
| Pyruvate | Agilent Technologies | 103578-100 | 1 mM |
| Seahorse XF Calibrant Solution | Agilent Technologies | 100840-000 |  |
| Seahorse XFp Real-Time ATP Rate Assay | Agilent Technologies | 103591-100 |  |
| Transferrin from Human Serum, Alexa Fluor™ 647 Conjugate | Invitrogen | T23366 | Sterile water<br>2 mM sodium azide<br>25 µg/mL |
| Magnesium Chloride | Caledon Laboratory Chemicals | 4720-1-70 | 500 mg/mL sterile water (100x)<br>1x in HBSS |
| CCCP | Sigma-Aldrich | 215911 | 90% Ethanol<br>10 µM |
| MitoTracker™ Red 580 | Molecular Probes | M-22425 | 100% DMSO<br>250 nM |
| TMRE | Invitrogen | T669 | 50% DMSO<br>30 nM |
| Seahorse XF HS Mini Analyzer | Agilent Technologies | S7852A | Cloud-based Seahorse Analytics |
| CytoSMART – Corning Cell Counter | Corning | 6749 | v.3 |
| TiEclipse inverted A1 confocal microscope | Nikon |  | NIS Elements AR 5.30.05v 64-bit<br>NIS Elements JOBS |
| Tokai Hit incubation system | Tokai Hit USA Inc. | INUBG2AT FP-WSKM |  |
| CellProfiler™ | Broad Institute | https://doi.org/10.5683/SP3/VII0UC | Version 4.2.5 |
| KNIME® | KNIME | https://doi.org/10.5683/SP3/VII0UC | 4.5.1v |
| GraphPad Prism | Dotmatics |  | Version 10.4.0 |

\*Gifted by Dr. Zhixing Chen from Peking University, China

**Table 3.** Absolute values, standard deviations, and variances of experimental results.

| **Experimental Condition** | **Reference Figure** | **Cell Type** | **Absolute Mean** | **Standard Deviation** | **Variance** |
| --- | --- | --- | --- | --- | --- |
| MitoTracker™ Red Intensity | Figure 3 A | Control | 0.2548 | 0.0843 | 0.0071 |
| PKmito™ Dye Red | Figure 3 A | Control | 0.8252 | 0.0437 | 0.0019 |
| MitoTracker™ Deep Red Intensity | Figure 3 B | Control | 0.0237 | 0.0079 | 0.0001 |
| PKmito™ Dye Red | Figure 3 B | Control | 0.4621 | 0.1106 | 0.0122 |
| CCCP Morphology - Eccentricity: No Treatment | Figure 3 C | Control | 0.7021 | 0.1370 | 0.0188 |
| CCCP Morphology - Eccentricity: Vehicle | Figure 3 C | Control | 0.7166 | 0.1308 | 0.0171 |
| CCCP Morphology - Eccentricity: CCCP | Figure 3 C | Control | 0.6173 | 0.1553 | 0.0241 |
| CCCP Morphology – Area: No Treatment | Figure 3 D | Control | 0.8253 | 0.3898 | 0.1519 |
| CCCP Morphology – Area: Vehicle | Figure 3 D | Control | 0.7356 | 0.3602 | 0.1297 |
| CCCP Morphology – Area: CCCP | Figure 3 D | Control | 1.066 | 0.6439 | 0.4146 |
| CCCP Morphology - Compactness: No Treatment | Figure 3 E | Control | 1.144 | 0.6313 | 0.3985 |
| CCCP Morphology - Compactness: Vehicle | Figure 3 E | Control | 1.310 | 0.6503 | 0.4229 |
| CCCP Morphology - Compactness: CCCP | Figure 3 E | Control | 1.486 | 0.9842 | 0.9686 |
| Microscope Comparison – Eccentricity: Widefield | Figure 4 A | Control | 0.9525 | 0.0375 | 0.0014 |
| Microscope Comparison – Eccentricity: | Figure 4 A | Control | 0.9206 | 0.0409 | 0.0017 |
| Confocal |  |  |  |  |  |
| Microscope Comparison – Confocal: Confocal | Figure 4 B | Control | 2.131 | 1.381 | 1.907 |
| Microscope Comparison – Compactness: Confocal | Figure 4 B | Control | 1.505 | 0.5719 | 0.3271 |
| Objective Comparison – Eccentricity: 20x | Figure 4 C | Control | 0.9206 | 0.0409 | 0.0017 |
| Objective Comparison – Eccentricity: 60x | Figure 4 C | Control | 0.9153 | 0.0234 | 0.0005 |
| Objective Comparison – Compactness: 20x | Figure 4 D | Control | 1.505 | 0.5719 | 0.3271 |
| Objective Comparison – Compactness: 60x | Figure 4 D | Control | 2.535 | 0.4309 | 0.1857 |
| Live vs Fixed Comparison – Eccentricity: Live | Figure 4 E | Control | 0.9153 | 0.0234 | 0.0005 |
| Live vs Fixed Comparison – Eccentricity: Methanol | Figure 4 E | Control | 0.8770 | 0.0210 | 0.0004 |
| Live vs Fixed Comparison – Eccentricity: PFA | Figure 4 E | Control | 0.8719 | 0.0246 | 0.0006 |
| Live vs Fixed Comparison – Compactness: Live | Figure 4 F | Control | 2.535 | 0.4309 | 0.1857 |
| Live vs Fixed Comparison – Compactness: | Figure 4 F | Control | 3.051 | 0.6764 | 0.4575 |
| Methanol |  |  |  |  |  |
| Live vs Fixed Comparison – Compactness: PFA | Figure 4 F | Control | 3.377 | 0.8315 | 0.6914 |
| Basal Morphology: Eccentricity | Figure 5 A | Control | 0.9049 | 0.0301 | 0.0009 |
| Basal Morphology: Eccentricity | Figure 5 A | HD | 0.8881 | 0.0318 | 0.0010 |
| Basal Morphology: Compactness | Figure 5 B | Control | 2.454 | 0.4735 | 0.2242 |
| Basal Morphology: Compactness | Figure 5 B | HD | 2.078 | 0.3768 | 0.1420 |
| Drp1 Intensity | Figure 5 E | Control | 0.0565 | 0.0167 | 0.0003 |
| Drp1 Intensity | Figure 5 E | HD | 0.0725 | 0.0248 | 0.0006 |
| glycoATP Production Rate | Figure 5 F | Control | 74.78 | 9.402 | 88.39 |
| glycoATP Production Rate | Figure 5 F | HD | 61.11 | 9.677 | 93.64 |
| mitoATP Production Rate | Figure 5 F | Control | 150.8 | 12.63 | 159.5 |
| mitoATP Production Rate | Figure 5 F | HD | 60.65 | 15.16 | 229.8 |
| Compensatory Glycolysis | Figure 5 F | Control | 323.7 | 23.51 | 552.7 |
| Compensatory Glycolysis | Figure 5 F | HD | 224.9 | 27.98 | 782.9 |
| Membrane Potential – PKMR: Vehicle | Figure 5 G | Control | 0.2447 | 0.0754 | 0.0057 |
| Membrane Potential – PKMR: Vehicle | Figure 5 G | HD | 0.2377 | 0.0777 | 0.0060 |
| Membrane Potential – PKMR: CCCP | Figure 5 G | Control | 0.2639 | 0.0547 | 0.0030 |
| Membrane Potential – PKMR:<br>CCCP | Figure 5 G | HD | 0.2562 | 0.0522 | 0.0027 |
| Membrane Potential – TMRE:<br>Vehicle | Figure 5 H | Control | 0.0601 | 0.0202 | 0.0004 |
| Membrane Potential – TMRE:<br>Vehicle | Figure 5 H | HD | 0.0669 | 0.0216 | 0.0005 |
| Membrane Potential – TMRE:<br>CCCP | Figure 5 H | Control | 0.0349 | 0.0113 | 0.0001 |
| Membrane Potential – TMRE:<br>CCCP | Figure 5 H | HD | 0.0347 | 0.0127 | 0.0002 |
| Membrane Potential – MTR:<br>Vehicle | Figure 5 I | Control | 0.0275 | 0.0079 | 0.0001 |
| Membrane Potential – MTR:<br>Vehicle | Figure 5 I | HD | 0.0271 | 0.0081 | 0.0001 |
| Membrane Potential – MTR:<br>CCCP | Figure 5 I | Control | 0.0286 | 0.0072 | 0.0001 |
| Membrane Potential – MTR:<br>CCCP | Figure 5 I | HD | 0.0281 | 0.0074 | 0.0001 |
| Transferrin Intensity | Figure 5 J | Control | 0.0405 | 0.0082 | 0.0001 |
| Transferrin Intensity | Figure 5 J | HD | 0.0377 | 0.0116 | 0.0001 |

### Protocol optimization using control fibroblasts demonstrates the sensitivity of the mitochondrial morphology assessment system

A graphical representation of the mitochondrial morphology assessment system (MMAS) is presented in Figure 1. The fluorescence intensity of PKmito^TM^ Dye Red (PKMR) and Deep Red (PKMDR; Table 2) was compared with the popularized MitoTracker^TM^ Red (MTR) and Deep Red (MTDR) under the same conditions (Figures 3 A and B). Each stain was permitted to incubate at the same concentration for the same time before being replaced with imaging media. Both PKMR and PKMDR have significantly higher fluorescence intensities than their MitoTracker^TM^ counterparts (p <0.0001), with a median intensity almost three times brighter. Brighter fluorescence allows for the minimization of laser power, thereby reducing phototoxicity and preserving mitochondrial health.^27, 28^ In later experiments, treatment with PKMDR was preferred because it is less toxic to cells during high exposure to lasers.^29^ The far-red spectrum has a longer wavelength (640 nm) and, therefore, less energy to produce damage to cells when imaging.^30^

**Figure 1.**
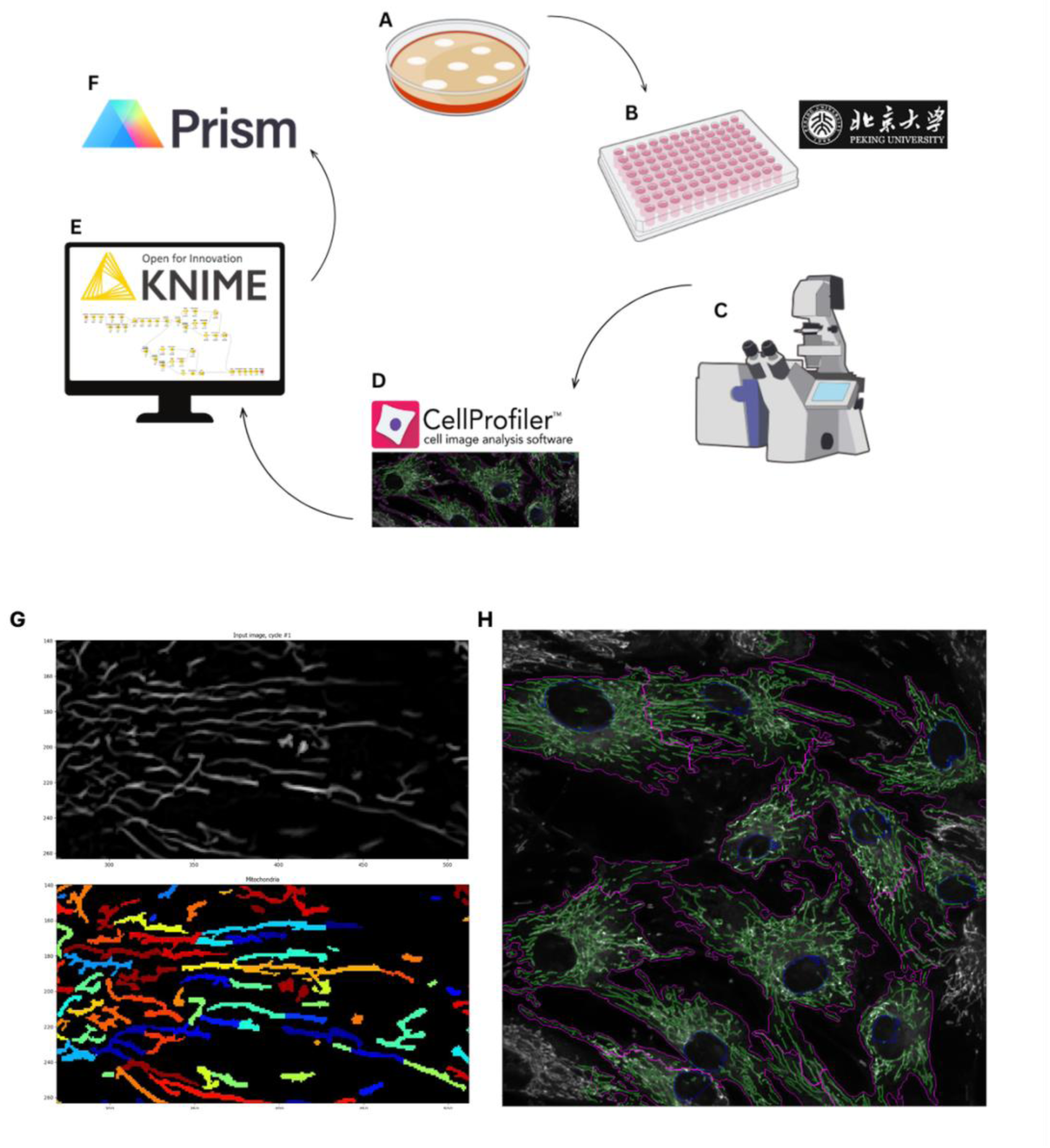
Graphical overview of the Mitochondrial Morphology Assessment System (MMAS). **(A)** Cells are seeded into imaging plates. **(B)** The new mitochondrial probe, PKmito^TM^ Dye, from the Chen Lab at Peking University and NucBlue^TM^ are used to stain cells for **(C)** confocal imaging (60x oil-immersion) using robotic stage and incubation control. **(D)** Exported images are uploaded into CellProfiler™ for identification and segmentation, where the purple outline represents the cell body, and the green outlines the mitochondria. **(E)** The resulting data sets are uploaded into KNIME^®^, which mines through the tens of thousands of rows of data and turns it into a document that can be directly plotted and statistically analyzed using GraphPad Prism. Illustrations in **(A-C)** are from NIAID NIH BIOART and the Figure containing **(A-F)** was generated by Amy Klarer. **(F-G)** Representative images from CellProfiler^TM^-based identification and segmentation of mitochondria. **(F)** An example of mitochondrial structures before (top) and after (bottom) segmentation, where distinct mitochondrial filaments are pseudo-coloured. **(G)** Overlayed outlines on the original confocal image depict nuclei (blue), cell body (purple), and mitochondria (green) and are exported during analysis.

**Figure 2:**
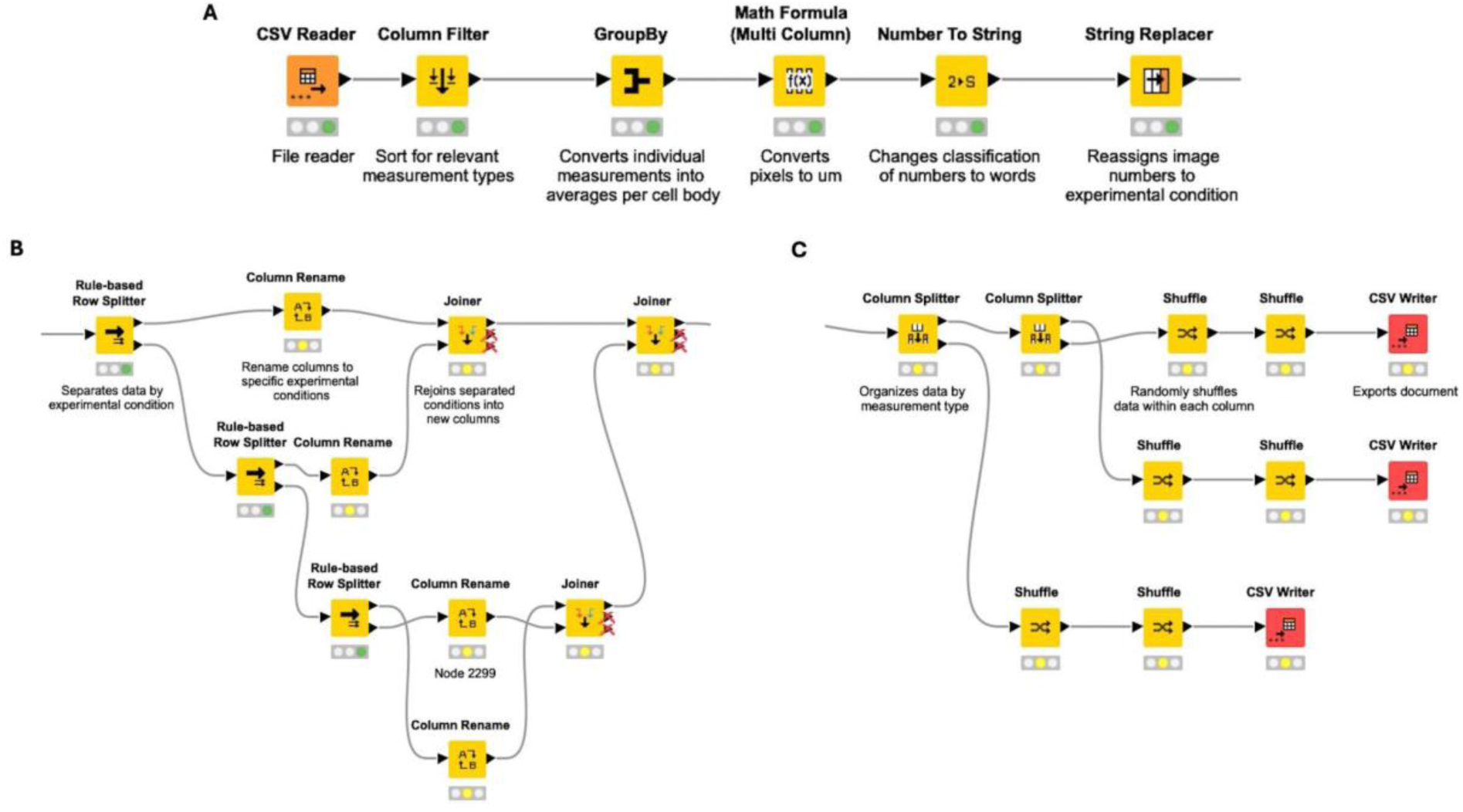
Representative images of the KNIME^®^ workflow for processing CellProfiler^TM^ data. **(A)** The pipeline uploads raw CSV data, filters by relevant measurements, calculates averages per cell body, and converts pixel-based measurements to micrometers. **(B)** Rule-based sorting and column renaming organize data by experimental condition. **(C)** Data is extracted into three documents based on the desired measurement type, randomized, and exported for further statistical analysis.

**Figure 3:**
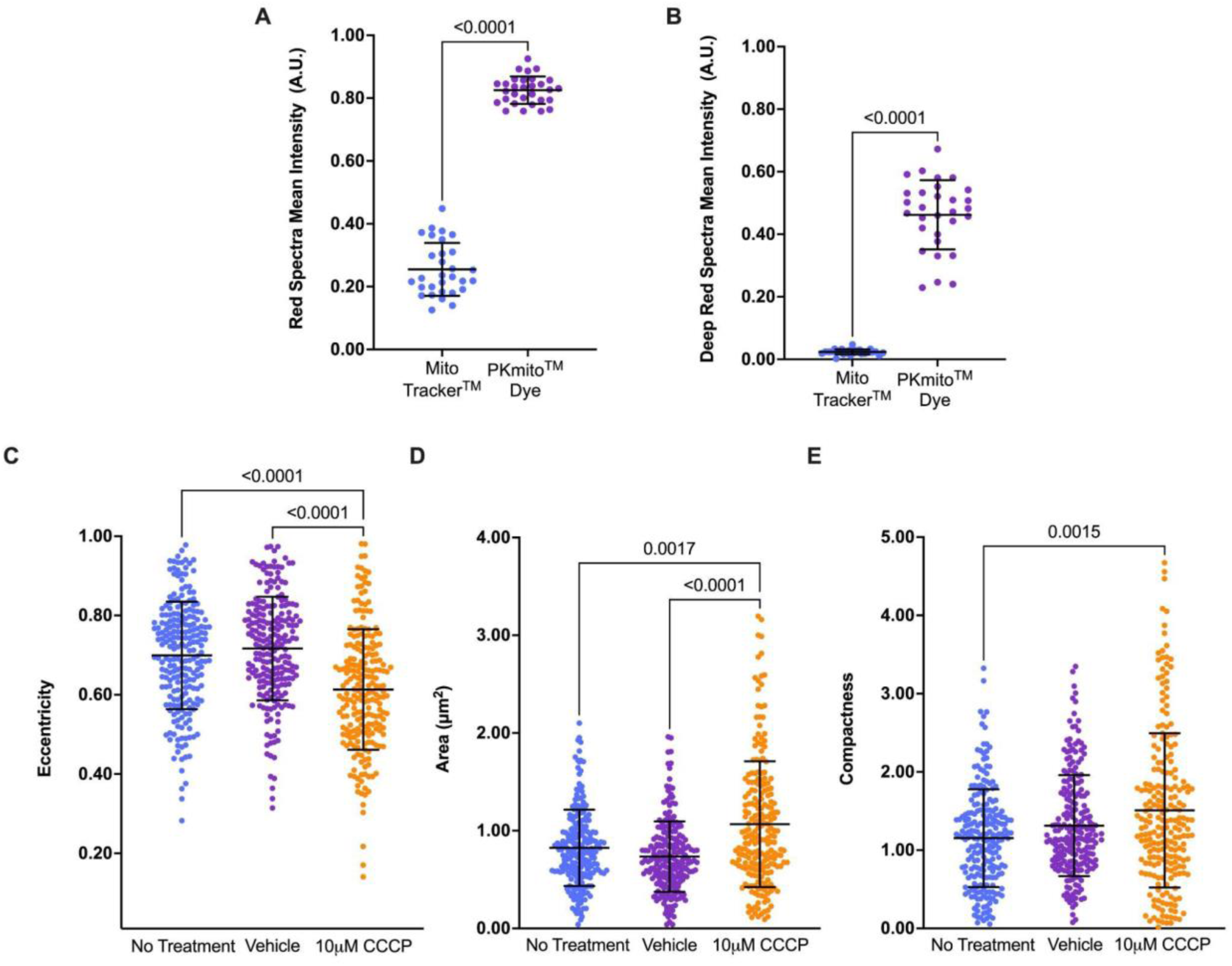
Protocol optimization of mitochondrial staining and validation of the mitochondrial morphology assessment system in control fibroblasts. Intensity comparison of PKmito^TM^ Dye and MitoTracker™ staining in red **(A)** or deep red **(B)** spectra. PKmito^TM^ Dyes exhibit significantly brighter fluorescence intensity than MitoTracker™ (Mann-Whitney test), allowing for improved mitochondrial visualization. Data represent the mean fluorescence intensity per cell body (N = 3, n = 30). **(C-E)** Demonstrate the validity of MMAS sensitivity using CCCP-induced mitochondrial stress. **(C)** The eccentricity significantly decreases following CCCP treatment, indicating increased mitochondrial circularity. **(D)** The mitochondrial area increases under CCCP treatment compared to no treatment and compared to the vehicle, which is consistent with mitochondrial swelling. **(E)** Compactness measurements increase slightly after treatment with CCCP, suggesting structural changes associated with depolarization. All data in C-E are represented as scatter plots +/- SD with N=3, n=223 where n=average mitochondrial measurement per cell body and N=number of experimental replicates. Statistical significance was determined using the Kruskal-Wallis and Dunn’s multiple comparisons test.

Our CellProfiler^TM^ pipeline was adapted from the protocol outlined by Rees et al. (2020), who used a fixed neuronal mouse model of Parkinson’s Disease.^31^ Images in their study were obtained using 40x widefield microscopy.^31^ Due to the intrinsic limitations of widefield microscopy, uneven illumination across the images must be calculated and corrected, which is time-consuming.^31–33^ Identification and segmentation of the mitochondrion has to be rigorous and highly accurate because of the increased complexity of neuronal morphology.^34, 35^ In addition, fixation can introduce artifacts or permanently alter cellular structures.^36, 37^ The introduction of any change in experimental design requires some degree of customization of the image analysis protocol.

The morphometric measurements examined in this study include eccentricity, area, and compactness. Eccentricity (E) quantifies the elongation of the mitochondria, calculated as the ratio of half the minor axis length to the major axis length. A value of 1 represents linearity, and 0 represents a circle. Area (A) measures the space occupied by a mitochondrion, providing insight into changes in size that could result from fragmentation or fusion. Compactness (C) is given as the perimeter divided by 4πA. A perfect circle has a C value of 1, whereas higher numbers reflect irregularities in the object’s shape and size, characterizing a fragmented mitochondrion shape.^38^ Since a given mitochondrion may maintain its total area while changing shape dramatically, eccentricity and compactness are more informative indices for understanding structural dynamics.

We tested the sensitivity of the MMAS by inducing mitochondrial damage using carbonyl cyanide 3-chlorophenylhydrazone (CCCP; Figures 3 C-E). CCCP is a protonophore, depolarizing the mitochondrial membrane by disrupting the proton gradient.^39^ CCCP-treated mitochondria showed significantly decreased eccentricity (p <0.0001) compared to the vehicle and negative control. With regard to their area, CCCP treatment enlarged mitochondria relative to the vehicle (p <0.0001) and negative (p = 0.0017) controls. Though more subtly, the same trend was demonstrated with compactness (vehicle = n.s.; negative = 0.0024), confirming that the MMAS can detect mitochondrial stress induced by cell treatment with CCCP. Presumably, a collapse of the proton gradient and an osmotic imbalance following membrane depolarization caused swelling of the mitochondria so that they become larger and rounder.^40^

### Technical optimization of microscopy methods using control fibroblasts reveals the importance of high-resolution and live-cell imaging

Next, we aimed to reduce excess photobleaching and phototoxicity by assessing the ideal imaging conditions. Both widefield and confocal microscopy detect changes in mitochondrial morphology (Figures 4 A, B). However, confocal imaging produced lower values for eccentricity (p <0.0001) and compactness (p = 0.0007) than widefield. Although it is not possible to independently determine the accuracy of one method over the other, we can compare the spread in the data, which is indicative of the measurement precision. Both types of microscopes produced eccentricity measurements with similar variance, suggesting one technology has no advantage over another. There is, however, a large discrepancy in the data variance (*s^2^*) when assessing compactness, with widefield microscopy (*s^2^* = 1.907) having a much broader distribution than confocal (*s^2^* = 0.3271). As a consequence, confocal microscopy was preferred over widefield modality in subsequent experiments.

**Figure 4:**
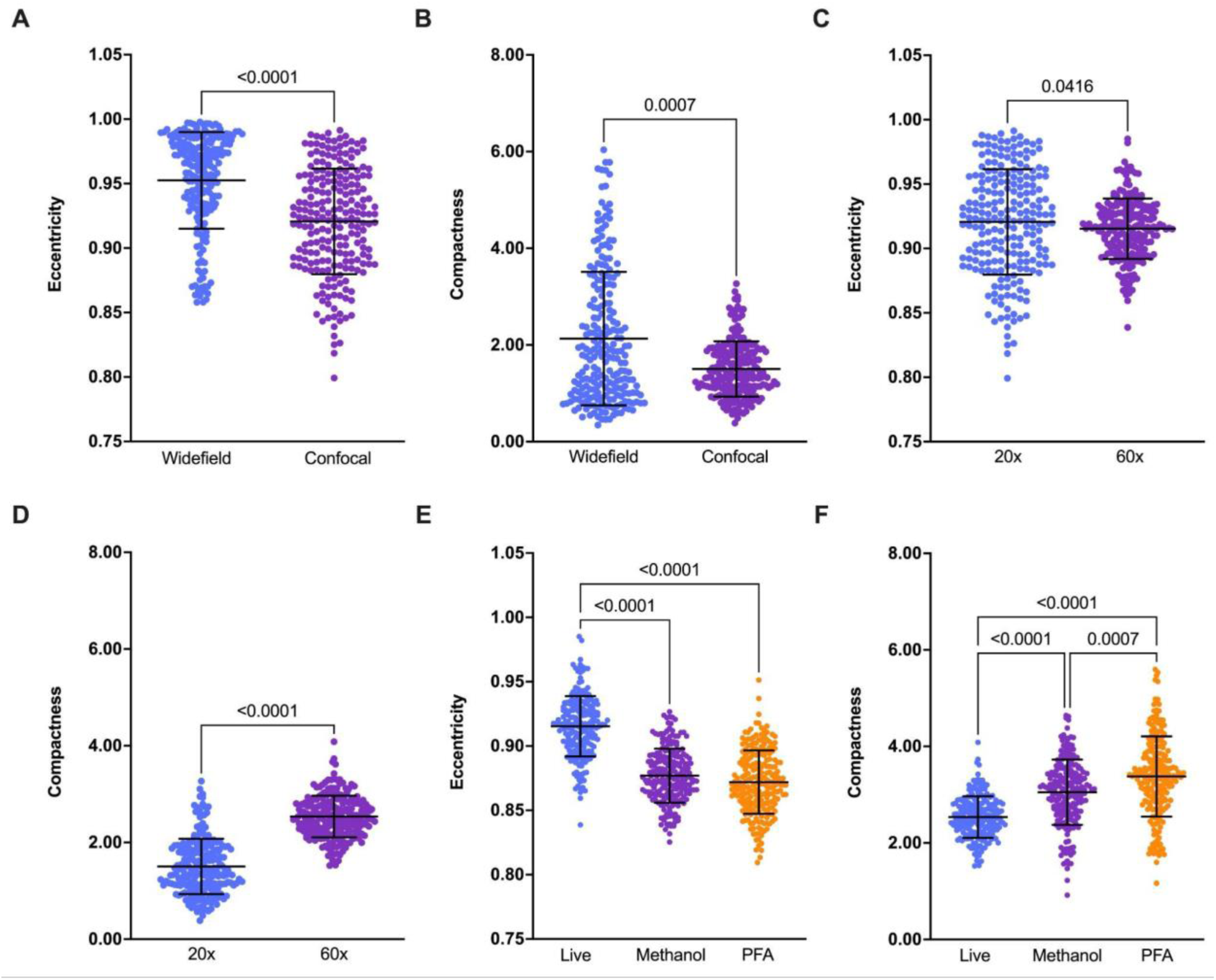
Technical optimization of microscopy techniques for analyzing mitochondrial morphology in control cells. **(A, B)** Comparison of widefield and confocal microscopy for imaging eccentricity **(A)** and compactness **(B)**. Confocal imaging resulted in significantly lower eccentricity and compactness values than widefield. **(C, D)** Comparison of 20x air and 60x oil-immersion objectives for eccentricity **(C)** and compactness **(D)**. Higher magnification (60x) resulted in lower eccentricity and higher compactness. **(E, F)** differences in live-cells and cells fixed via two fixation methods: 100% (v/v) methanol and 4% paraformaldehyde (PFA) for eccentricity **(E)** and compactness **(F)**. Fixed samples have lower eccentricity and increased compactness compared to live-cells. PFA significantly increases compactness compared to methanol fixation. Data are represented as scatter plots +/- SD with N=3, n=223 where n=average mitochondrial measurement per cell body and N=number of experimental replicates. Statistical significance was determined using the Mann-Whitney **(A-D)** or Kruskal-Wallis test with Dunn’s multiple comparisons test **(E, F)**.

#Figures 3 C and D evaluate the influence of objective magnification (i.e., 20x air versus 60x oil-immersion). Measurements of mitochondrial eccentricity were relatively consistent in the mean between the two; however, the variance for the 20x objective was twice as large (*s^2^* = 0.0017) as that for 60x (*s^2^* = 0.0005). When assessing compactness, mitochondria measured significantly higher (p <0.0001) using the 60x objective as compared to 20x, despite more similar variances (*s^2^* = 0.3271 and 0.1857, respectively). These results support that the increased resolution provided by the 60x oil-immersion objective is more desirable for morphological quantification due to its increased precision.

In a final step towards optimizing the MMAS, we compared live- and fixed-cell imaging. Fixed-cell imaging involved one of two treatments: 15 minutes of methanol or 20 minutes with 4% PFA and permeabilization. Figures 3 E and F show that mitochondria from live-cells have a significantly higher eccentricity than fixed samples (p <0.0001), indicating that fixation alters mitochondrial morphology. No difference in variance was found between live, methanol-fixed, or PFA-fixed trials. In comparison, the variance was found to increase for measurements of compactness in live-cells (*s^2^* = 0.1857) and those treated with methanol (*s^2^* = 0.4575) or PFA (*s^2^* = 0.6914). Mean compactness increased significantly between live- and fixed-cells (p <0.0001) and increased again between methanol and PFA (p = 0.0007). Though it remains impossible to determine which absolute mean is more accurate, a clear advantage is associated with the high precision demonstrated for live-cell imaging.

### Mitochondrial dysfunction is quantified using the MMAS and verified with orthogonal assays

HD cells were compared with those from a wildtype spousal control (Table 1) to test the potential of the MMAS as a tool for distinguishing between healthy and diseased mitochondria. As shown in Figures 5 A and B, HD mitochondria were significantly rounder (p <0.0001) and more compact (p <0.0001) than the control, consistent with a fragmented phenotype typically associated with fission.^41, 42^ The increased fragmentation of HD mitochondria is also evident to the human eye in Figures 5 C and D, further confirming the efficacy of the MMAS.

**Figure 5.**
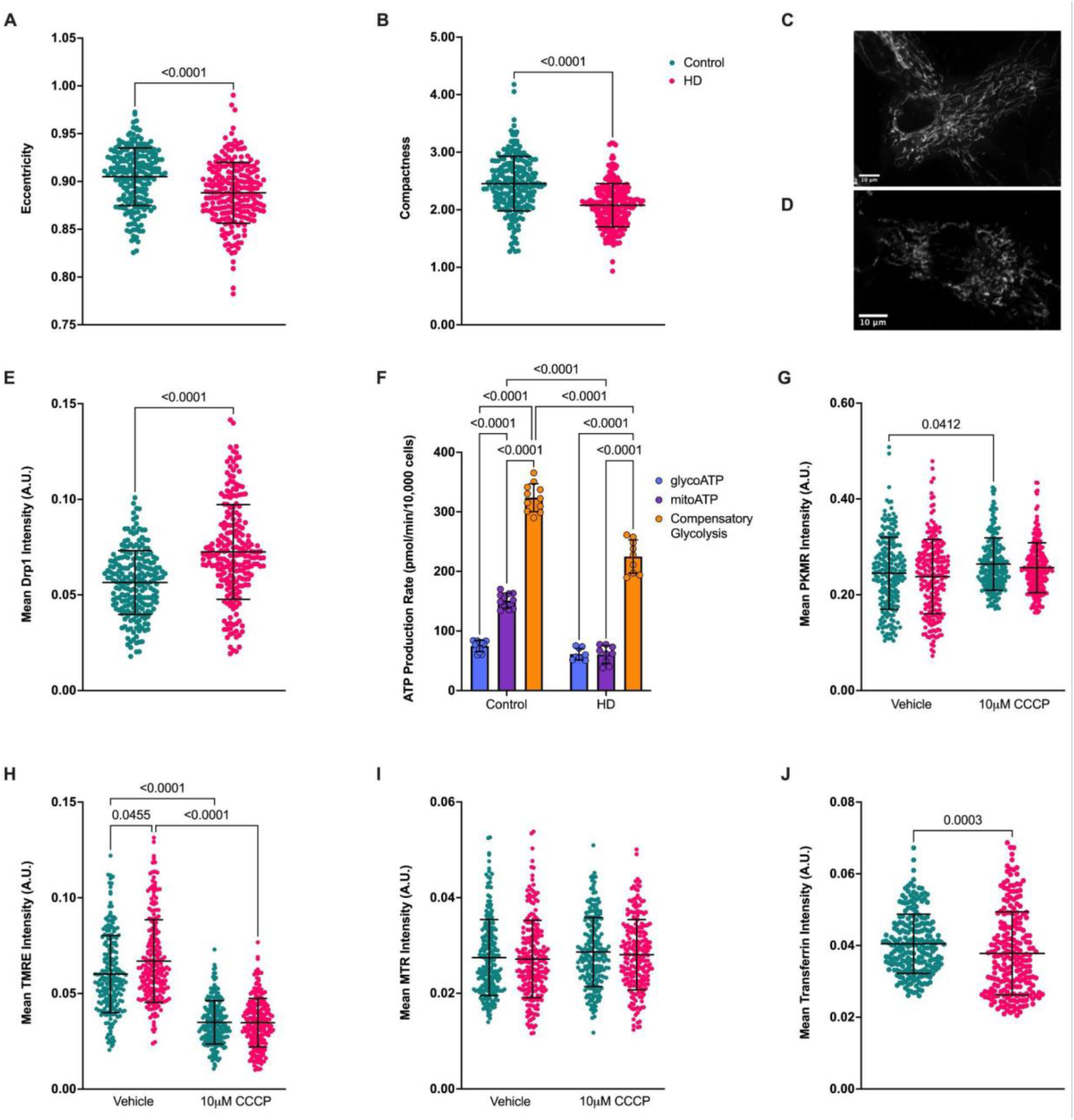
Mitochondrial morphology, bioenergetics, and membrane potential in healthy and HD fibroblasts. **A** and **B** show the quantification of eccentricity **(A)** and compactness **(B)** in control and HD cells. HD mitochondria are significantly rounder and more compact than the control (N=3, n=223; Mann-Whitney test). Representative images of mitochondrial networks in healthy **(C)** and HD **(D)** cells, showing fragmentation in HD (scale bar = 10 μm). **(E)** The mean fluorescence intensity of the key fission regulator, Drp1, demonstrated significantly increased levels in HD (N=3, n=223; Mann-Whitney test). **(F)** ATP production rate in glycolytic, mitochondrial, or compensatory glycolysis between healthy (N=4, n=11) and HD (N=3, n=9) cells. Control cells rely more on oxidative phosphorylation for basal respiration than HD (2way ANOVA with Šídák’s multiple comparisons test). **(G-I)** Mitochondrial membrane potential using various fluorescent dyes (N=6, n=223). **(G)** PKmito^TM^ Dye Red intensity, **(H)** TMRE intensity, and **(I)** MitoTracker^TM^ Red intensity in vehicle-treated and CCCP-treated cells. CCCP depolarization is only effective in TMRE-stained cells. In comparison to the control, HD cells demonstrate increased membrane potential (Kruskal-Wallis test with Dunn’s multiple comparisons test). **(J)** Mean fluorescence intensity of transferrin, depicting a significant decrease in HD cells compared to control (Mann-Whitney test). Data are represented as scatter plots +/- SD with N=number of experimental replicates and n=average mitochondrial measurement per cell body **(A, B, E, G-J)** or as the average ATP production rate per well **(F)**.

To determine if this fragmentation results from upregulated fission, levels of the mitochondrial fission protein Drp1 were quantified using immunofluorescence. As demonstrated in Figure 5 E, the relative fluorescence of Drp1 in HD is significantly higher than the control (p <0.0001), supporting the idea that the fragmented mitochondria are a result of increased fission.

We next asked whether increased fission in HD fibroblasts corresponded with changes in energy production since several studies have linked fission to glycolytic reprogramming via oxidative stress.^43, 44^ We used the Seahorse ATP Rate Assay to measure glycolytic (glycoATP), mitochondrial (mitoATP), and compensatory glycolysis. As shown in Figure 5 G, control lines primarily relied on oxidative phosphorylation (OXPHOS) rather than glycolysis (p < 0.0001), whereas HD fibroblasts did not prefer one method over the other. Both cell lines produced a similar amount of ATP from glycolysis, but control cells produced significantly more ATP through OXPHOS (p <0.0001). When forced to switch from OXPHOS to glycolysis (compensatory glycolysis), HD fibroblasts were considerably less responsive (p <0.0001). These analyses indicate that HD cells are metabolically less active and produce less energy than the wildtype cells, as anticipated with increased fission.^45^

Since HD mitochondria have proven to be dysfunctional, we wanted to determine if these abnormalities were reflected in mitochondrial membrane potential. Assessing mitochondrial membrane potential by quantifying fluorescence intensity allows for the comparison of traditional methods using tetramethylrhodamine, ethyl ester (TMRE) to the less popular MitoTracker^TM^ Red (MTR) and the new PKMR.^46^ As depicted in Figures 5 G-I, CCCP-induced membrane depolarization was detected with TMRE (p <0.0001), but not with PKMR or MTR. Since TMRE was the only probe that reacted to CCCP treatment, only experiments in Figure 5 I could be accurately interpreted. In contrast to the literature, HD cells demonstrate higher membrane potential than control (p = 0.0455).^23, 47–50^

Mitochondria are the primary regulators of cellular iron.^1^ Failure in managing high iron levels results in activation of ferroptosis, a nonapoptotic programmed cell death.^23, 51–53^ Ferroptosis results in mitochondrial dysfunction, such as fragmentation via Drp1 activation.^23, 54–56^ HD patients have demonstrated increased mitochondrial iron uptake and decreased utilization. This results in iron accumulation, which is thought to contribute to mitochondrial dysfunction.^54, 55^ Therefore, we assessed the involvement of an iron-binding glycoprotein called transferrin, which is responsible for transporting dietary iron across the blood-brain barrier.^57–61^ Human transferrin conjugated to Alexa Fluor^TM^ 647 for fluorescent intensity measurements was quantified. We saw a significant reduction of transferrin relative intensity in HD compared to the wildtype control (Figure 5 E, p = 0.0002). Less transferrin concentration suggests reduced iron uptake, possibly contributing to bioenergetic defects in HD.^52, 54, 55^

## Discussion

Here, we developed a novel system for quantifying specific morphometric measurements of mitochondria and other cellular components. This analysis technique uses the newly developed mitochondrial probe, PKmito^TM^ Dye, in live-cells with semi-automated, high content quantification techniques. The preliminary results demonstrate a significantly higher fluorescence capacity of the PKmito^TM^ Dye compared to MitoTracker^TM^ equivalents. The brighter fluorescence intensity of a probe allows for the reduction of laser power, thereby minimizing phototoxicity and preserving mitochondrial function. Microscopy and image quantification were partially automated using NIS Elements JOBS, CellProfiler^TM^, and KNIME^®^ to create a high content morphology assessment. Quantifying morphology in a non-biased manner using state-of-the-art, but open-access and user-friendly technology and methods will advance our fundamental understanding of structural mitochondrial dynamics in the context of various disease applications.

Treatment of healthy fibroblasts with the oxidative phosphorylation uncoupler, CCCP, induces mitochondrial fragmentation into small, globular structures, thereby providing a valuable tool for assessing the performance of the MMAS.^62^ CCCP treatment in this study resulted in a significant decrease in mitochondrial eccentricity, while the area was observed to increase. The larger and rounder morphology suggests a collapse of the proton gradient and an osmotic imbalance following membrane depolarization, causing swelling of the mitochondria.^40, 63^ These findings validate the MMAS for accurate, high content assessment of mitochondrial morphology.

An important technical goal for this study involved establishing the resolution requirements for the microscope. Notably, confocal microscopy captures a single plane in focus using a pinhole, whereas widefield captures images of the entire sample, including anything out of focus.^32^ As expected, both the widefield and confocal microscopes successfully detected change in the morphometric indices targeted; however, confocal imaging demonstrated higher precision, particularly when assessing compactness. With regard to a comparison of 20x air with 60x oil-immersion magnification, both objectives provided consistent results for mitochondrial eccentricity. At 60x, the variability of eccentricity and compactness decreased, while the absolute measurements of compactness were higher. These results were expected since the higher numerical aperture of the 60x objective allows for enhanced resolution of small morphological features, like mitochondria, and is especially important for detecting subtle changes.^64^ These findings suggest that higher resolution provides precise and consistent results.

Previous mitochondrial imaging studies have largely relied on fixed-cell samples, especially for techniques involving electron microscopy or immunofluorescence.^24, 65, 66^ Although the results derived from fixed-cells generally show preserved mitochondrial structure, an increased variance of eccentricity and compactness suggests the irreversible alteration of morphology or organization.^36, 37^ Artifacts interfere with reliable quantification; therefore, live-cell imaging is preferred. Live-cell imaging allows for the definitive assessment of mitochondria in their physiological environment, as required for understanding spatiotemporal cellular changes in real-time.

Using the MMAS to detect morphological differences between healthy and HD patient fibroblasts further demonstrates its potential as an analytical tool. HD mitochondria were found in this study to be rounder and more compact, consistent with a fragmented phenotype and associated with increased fission. Although statistically significant, the difference in absolute magnitude is relatively small. The rate of disease progression provides a possible explanation for this outcome: HD is late-onset, suggesting that relatively inconsequential abnormalities accumulate throughout a lifetime until substantial enough to produce symptoms.^12, 22^ Thus, this methodology may be useful in the context of late onset genetic diseases with subtle phenotypic affects that are nonetheless important to disease.

Previous studies have demonstrated increased expression of the fission genes *Drp1* and *Fis1*.^67–69^ We aimed to confirm that the fragmentation in HD mitochondria is associated with dysregulated Drp1 and further validate the phenotypes represented by the MMAS. Immunofluorescence results demonstrate significantly increased signal intensity in HD cells, substantiating the idea of upregulated fission. Upregulated Drp1 activity is demonstrated with reduced ATP production and increased reactive oxygen species generation, worsening mitochondrial stress and dysfunction.^70–72^

Fragmented mitochondria are often less efficient at oxidative phosphorylation (OXPHOS).^70^ Our results reinforce the link between fission and impaired energy production. Healthy cells primarily rely on OXPHOS for ATP production, whereas HD cells have no preference for OXPHOS over glycolysis. This lack of preference is as expected from the literature, where HD cells demonstrate abnormalities within specific complexes of the electron transport chain (ETC).^73, 74^ However, we anticipated an increase in reliance on glycolysis for ATP production, a hypothesis not supported by these results.^75^ HD cells are likely relying on a compensatory shift in a primary production pathway, for example, to fatty acid oxidation.^73^

In contrast to the literature, human HD mitochondria had elevated membrane potential.^23, 47–50^ An explanation for this is a compensatory mechanism where the ETC attempts to increase proton pumping with complexes I, III, and IV to maintain ATP production under stress, such as that from HD’s dysfunctional OXPHOS complexes II, III, and IV.^74, 76^ Another possible mechanism suggests hyperpolarization of mitochondria indicates stress and dysfunction rather than efficient biogenesis, supported by previous studies which found hyperpolarization before mitochondrial failure in neurodegenerative diseases.^77, 78^

Transferrin levels were assessed to understand a possible mechanism behind the mitochondrial dysfunction in HD, given the established link between iron metabolism and mitochondrial health.^79, 80^ Consistent with the literature, HD cells demonstrated reduced iron uptake, suggesting iron overload.^52, 54, 55^ Since mitochondria are the primary regulators of cellular iron, which is needed for effective electron transfer in the ETC and other mitochondrial processes, dysregulated iron transport likely worsens mitochondrial dysfunction in HD.^48^ Future studies should explore whether restoring iron homeostasis using chelating therapies can mitigate mitochondrial abnormalities in HD cells.^81, 82^

The advancements in methodology presented in this study offer a partially automated workflow for a completely unbiased morphological assessment of mitochondria, using open-access and user-friendly technology that will contribute to the fundamental understanding of structural dynamics with varying applications. HD is not the only neurodegenerative disease that is affected by metabolic dysfunction. Parkinson’s Disease and Alzheimer’s Disease demonstrate similar abnormalities.^83, 84^ Expansion of our understanding of mitochondrial dynamics will deepen our insight into the underlying disease mechanisms and allow for more informed development of therapeutic approaches, such as restoring metabolic functioning to improve quality of life.^85^

Future studies should extend this analysis to isogenic models of HD in live human cells to rule out any environmental and genetic modifiers that could impact these results.^22, 86, 87^ Assessing phenotypes from various disease stages (pre-symptomatic to late disease) would be invaluable for understanding disease progression. Investigating the impact of culture conditions, like metabolite concentrations, would further aid in discovering disease mechanisms and areas of intervention by mimicking physiological and environmental stressors. In support of these new directions, we suggest that MMAS will be an invaluable preclinical tool for assessing drug efficacy in restoring mitochondrial function.

## Materials and Methods

For a complete list of reagents and chemicals with reconstitution and dilution information, see Table 2. The general workflow for assessing mitochondrial morphology (Figure 1) involves seeding fibroblasts into a 96-well plate and growing until 90% confluent. Cells are incubated with the mitochondrial probe PKmito^TM^ Dye Deep Red (PKMDR), followed by the nuclear stain NucBlue^TM^.^9^ The plate is imaged using confocal microscopy and automatic stage control. Microscope images are quantified for object shape and size using CellProfiler^TM^, and the resulting data is mined using KNIME^®^ (Figure 2). The output is statistically analyzed and plotted for visualization with GraphPad Prism.

### Cell Culture and Treatments

hTERT-immortalized patient fibroblasts (TruHD cells; Table 1) were labelled as either “control” or “HD”.^17^ Control cells from a female donor contained CAG repeat lengths of 21 (non-pathogenic) and 18 (non-pathogenic), while HD cells from a male donor contained CAG repeat lengths of 43 (pathogenic) and 17 (non-pathogenic). HD cells were given a CAG-Age Product (CAP) score of 108, where 100-290 suggests early progression.^86^Both cell lines were authenticated using a PCR allele assay to confirm CAG repeat lengths and were regularly tested for mycoplasma contamination. Cells were cultured in MEM containing phenol-red and nonessential amino acids, supplemented with 15% fetal bovine serum (FBS) and a 1x concentration of GlutaMAX^TM^. Cells were incubated at 37°C with 5-8% oxygen and 5% carbon dioxide (CO_2_). Fibroblasts were seeded and grown until confluent in 96-well imaging plates for experimental use unless otherwise described.

### Mitochondrial Morphology

To objectively quantify morphological features like eccentricity, area, and compactness in mitochondria, fibroblasts were incubated with 250 nM PKMDR for 25 minutes, followed by 10 minutes of NucBlue^TM^ treatment (1 drop/mL media). Wells were washed with PBS and then replenished with imaging media: phenol-red free MEM supplemented with 15% FBS and 1x GlutaMAX^TM^ (from a 100x concentrated stock). The plates were imaged as described below using a 60x oil-immersion objective on a confocal microscope.

When assessing the effect of mitochondrial stress on morphology, cells were treated with 10 μM carbonyl cyanide m-chlorophenyl hydrazone (CCCP), ethanol, or media for 15 minutes post-staining. To quantify the impact of fixation on morphology, cells were either treated as outlined in the paragraph above or fixed with methanol or PFA. Fibroblasts fixed with cold methanol were incubated for 15 minutes at -20°C and then washed with cold PBS. Using 4% PFA, cells were incubated at room temperature for 20 minutes. PFA-treated cells were permeabilized with 20-minutes of 0.2% Triton^TM^ X-100 and then washed with PBS.

### Drp1 Immunofluorescence

Fibroblasts were fixed with 100% (v/v) cold methanol for 15 minutes at -20°C, then washed with cold PBS. Fixed cells were blocked for 15 minutes in blocking buffer (10% FBS diluted in PBS) before incubation with the primary anti-Drp1 antibody (1:100) for one hour. Cells were washed and then incubated for 15 minutes with the secondary Cy5® antibody (1:500). After washing again, the plate was incubated for 5 minutes with 0.2 μg/mL Hoechst before imaging with the 20x air objective on the confocal microscope.

### Seahorse XF Analysis

The Agilent Seahorse XF HS Mini Analyzer was used to quantify bioenergetics in fibroblasts. A CytoSMART - Corning Cell Counter v.3 was used to quantify cellular concentration. Cells were diluted to 5.0 x 10^4^ cells per well (50 μL) and seeded into wells B through G of an 8-well Seahorse PDL miniplate. In addition, 50 μL of MEM that did not contain cells were placed in wells A and H as a negative control. Moats around the wells were filled with 400 μL of sterile water and incubated overnight. The flux cartridge was hydrated using sterile water and incubated overnight in a non-CO_2_ 37°C incubator.

The next day, the Seahorse XF HS Mini Analyzer was allowed to equilibrate to 37°C. Media from the overnight culture was exchanged for XF assay media (DMEM with 10 mM glucose, 2 mM glutamine, and 1 mM pyruvate). Cells were monitored for adherence to the plate before and after this process. The miniplate was then placed in a non-CO_2_ incubator at 37°C for an hour with the cartridge. Sterile water in the cartridge was replaced with the pre-warmed calibrant and put back in the incubator for at least an hour before running the assay. The Seahorse ATP Rate Assay was run with increased concentrations of oligomycin (3 μM) and rotenone plus antimycin A (1 μM) due to the relative inactivity of the TruHD cells. In addition, 10x NucBlue^TM^ was added to the final injection port in conjunction with the last inhibitor.

The results were normalized to the total cell count per well to account for any differences in seeding density. Cells were imaged in PBS using the 10x objective lens on a confocal microscope using a custom 3D printed stage adaptor designed and printed by the Truant lab to lower the plate into the microscope’s dynamic focal range. Images of all wells were collected, including A and H to confirm the absence of cells. Once the data were analyzed, a calculation was completed to convert the number of nuclei in an image to the total number of cells per well, as depicted in Equation 1. The resulting concentrations were uploaded into the cloud-based Seahorse Analytics platform under the normalization setting.

**Equation 1**. The calculation used to validate the total number of cells per well in a Seahorse PDL Miniplate to normalize the results of an assay. Image width and length measurements are in pixels, and the conversion factor is in μm/pixel, which can be found in the “Image Properties” of the .nd2 files from the microscope. Final cell counts can then be uploaded into the Seahorse Analytics Platform for normalization. NF = normalization factor.

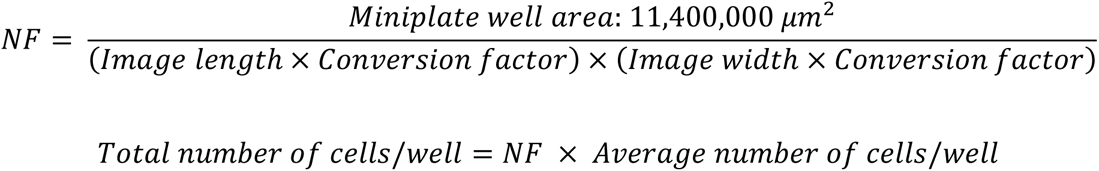

### Immunofluorescence Quantification of Transferrin Uptake

Assay buffer (HBSS supplemented with 1x calcium and magnesium (Ca^2+^ Mg^2+^), 20 mM Glucose, and 10% FBS) was divided into two; half was chilled on ice, and the other was warmed to 37°C. The plate of cells was chilled on ice for 10 minutes, after which the cells were washed with cold assay buffer. Conjugated human transferrin was diluted in pre-warmed assay buffer to 25 μg/mL and placed in each well for 20 minutes. Once completed, phenol red-free MEM supplemented with 20 mM glucose and 10% FBS was used to wash cells and then replenished for imaging. This plate was imaged with the 20x objective on a confocal microscope.

### Mitochondrial Membrane Potential

Confluent fibroblasts in 96-well plates were separated into six quadrants: no treatment, ethanol control, and 10 μM CCCP for staining with either 250 nM PKmito^TM^ Dye Red (PKMR), 250 nM MitoTracker^TM^ Red, or 30 nM Tetramethylrhodamine, Ethyl Ester, Perchlorate (TMRE). All mitochondrial probes were incubated for 30 minutes and then replaced with NucBlue^TM^ staining for 10 minutes; PBS washes were completed before replacing with the phenol-red free imaging media described in “Mitochondrial Morphology”. Once set up, the cells were treated with either imaging media, ethanol, or CCCP for 15 minutes. This plate was imaged with the 20x air objective on a confocal microscope.

### Microscopy and Image Acquisition

All experiments used either the 60x oil-immersion (NA = 1.4), the 20x air (NA = 0.75), or the 10x air (NA = 0.75) objective on a Nikon TiEclipse inverted A1 confocal microscope driven by NIS Elements AR 5.30.05v 64-bit acquisition software. Experiments using live-cells were imaged using a Tokai Hit incubation system adapted for the microscope with 37°C, 5% CO_2_, and humidity control. NIS Elements JOBS was used to automate the acquisition process to randomize the location of 7 images per well within a set of rectangular boundaries, eliminating any edge effect and potential bias. Cells were imaged using the DAPI (405 nm) and TRITC (560 nm) or Cy5 (640 nm) lasers with a 1024-pixel image area, 2.4-pixel dwell time, and a pinhole of 1.2.

### Data Analysis

Microscopy images were exported in binary as Tagged Image File Format (TIFF) independent images separated by channel. These exported 12-bit images were uploaded into CellProfiler^TM^ version 4.2.5 for identification and quantification (Figure 1 D). Our CellProfiler^TM^ pipeline can be accessed through McMaster Dataverse, a collection within Borealis, the Canadian Dataverse Repository, using https://doi.org/10.5683/SP3/VII0UC.26

The first module, “IdentifyPrimaryObjects,” identified and segmented nuclei based on the size, shape, and intensity of the stained nuclei. The example pipeline provided in Borealis was developed for fibroblasts imaged by confocal microscopy with a 60x oil-immersion objective. This module must be adapted if any of these parameters were changed. The accepted diameter (in pixels) should be adjusted first if the pipeline is able to identify nuclei, but the measurements are filtered out. Thresholding parameters like “method,” “smoothing scale,” and “correction factor” could be changed if there were problems correctly identifying objects.

This pipeline was created using a “global,” “minimum cross entropy” thresholding strategy. A global strategy calculates the threshold based on a single (average) value of an input image, whereas an adaptive strategy calculates this value for each pixel in the image. An adaptive strategy is slower, but accounts for high variation within the background. An automatically calculated threshold for distinguishing foreground from background was completed using minimum cross-entropy. This thresholding strategy divides pixel intensities into two classes and should be implemented when there is uneven illumination or subtle differences in object intensity.^88^ Improper segmentation of nuclei can be addressed by altering “methods to distinguish clumped objects,” “method to draw dividing lines..,” and “suppress local maxima that are closer than…”. Distinguishing clumped objects and drawing dividing lines could be completed by object intensity or shape. Distinguishing by shape was best used for separating uniformly round objects, and by intensity was best when the edges of the objects were dimmer than their centre—an additional option of propagation was available when drawing dividing lines in cases of cells with branching extensions, such as neurites. Suppressing local maxima was used when too many objects were merged (under-segmented) or split (over-segmented). The value, in pixels, should be approximately equivalent to the radius of the smallest identified object.^89^

Using “IdentifySecondaryObjects,” the software identified and segmented the cell body by propagating outward from the nuclei, relying on a ratio between the intensity of the mitochondrial stain at each point relative to its distance from the nuclei, called the regularization factor. This module relied on “Otsu,” “three class” thresholding rather than the minimum-cross entropy employed previously. Three-class thresholding was used because of the variation in the intensity of mitochondrial objects. Background, dim signal, and bright signal were identified, and the mid-intensity pixels were assigned as “foreground”.^90^ Otherwise, the same thresholding parameters as the previous module can be altered to enhance the accuracy of object identification. This step was tedious, and the segmentation was not necessarily exact for every experimental circumstance. In this study, priority was given to maintaining consistency between the experimental trials such that suitable comparisons could be drawn.

Mitochondrial features were then enhanced using “EnhanceOrSuppressFeatures” based on neurites (i.e., long, thin objects were assigned increased intensity values) so they may be classified based on their intensity and size. Another “IdentifyPrimaryObjects” module was added to identify and segment mitochondria. The thresholding method employed was the two-class version of Otsu, which is valuable for distinguishing two distinct bright and dim populations of intensity.^90, 91^ Otherwise, the same troubleshooting techniques previously mentioned can be applied. Mitochondria were then assigned a “parent” cell body associated with their XY location in the “RelateObjects” module. Measurements of the size, shape, and intensity of the nuclei, cell body, and mitochondria were then quantified using “MeasureObjectSizeShape” and “MeasureObjectIntensity”.

These data, along with the corresponding images containing overlays of the object outlines (“OverlayOutlines”), were exported for analysis from the CellProfiler^TM^ pipeline (Figures 1 F, G) using “SaveImages” and “ExportToSpreadsheet”. The overlain images allow the investigator to check for identification issues and provide independent visual confirmation of the segmentation. For example, images with skewed data were cross-referenced with the corresponding overlain images (Figure 1 G) and were marked for exclusion from the dataset when the segmentation was poor.

The exported datasets were subsequently uploaded into KNIME^®^ 4.5.1v for data restructuring, mining, and computational analyses (Figure 2; this workflow can be accessed through McMaster Dataverse, a collection within Borealis, the Canadian Dataverse Repository, using https://doi.org/10.5683/SP3/VII0UC).^26^ Figure 2 A demonstrates the workflow for eliminating irrelevant measurements or columns. The remaining mitochondrial measurements (eccentricity, area, and compactness) were aggregated by mean into their corresponding cell body to account for variation in cellular conditions. Measurements obtained in pixels, such as area, were converted into micrometres (squared) using a multi-column math formula from the calibration calculated by the microscope. Right-clicking an image in NIS Elements Viewer opens a window with image properties, where “Calibration” is found under “Image Field”. The calibration provided a value of 0.21 micrometers per pixel for optics with 60x oil-immersion.

Image numbers were converted to a string classification for manipulation (Figure 2 A). Changing the number to word classification allowed for relabelling well and image numbers with experimental conditions. For example, wells 1-3 and images 1-15 contained the control conditions (healthy cells, no treatment - NT). These images in the “string replacer” node would be denoted using regular expression as “[1-9]|1[0-5]” and then assigned the name “Control, NT”. Images previously marked for exclusion based on poor segmentation were filtered out during this step. If image 5 were marked for exclusion, the resulting expression would be “[1-4]|[6-9]|1[0-5]”. The number of “string replacer” nodes depends on the number of experimental conditions. An analysis containing two cell lines with a negative control, a positive control, and a treatment would have six replacer nodes.

Corresponding data were separated by experimental condition (Figure 2 B). In the previous example, data would be split initially by treatment and then by cell type. Columns were renamed based on these conditions, i.e. “Control, NT, Mean Area (μm^2^)”, before rejoining using a full outer join. The joined table was structured so that each measurement, experimental condition, and cell type were assigned individual columns rather than stacked in rows. These columns were split by measurement type (Figure 2 C) into separate tables and randomly shuffled. Shuffling the data within each column randomized the measurements from each image, well, and trial so that a sub-population could be sampled.

### Statistical Analysis

Due to the high content nature of these experiments, we used Slovin’s formula (Equation 2) to determine a suitable sample size (n) for analysis.^92^ Unless otherwise stated, n = 223 measurement values for each of <u>𝐸</u>_𝑖_, <u>𝐴</u>_𝑖_, and <u>𝐶</u>_𝑖_, where i =1… n cell bodies, were obtained for each of the three independent experiments. Since the data were already randomly shuffled in the final steps of the KNIME^®^ pipeline (Figure 1 I), the first 223 measurements from the exported file were used for plotting. Statistical analyses and graphical representations of each sample were completed using GraphPad Prism Version 10.4.0. Outliers were removed using the ROUT (Q = 1%) method. Normality distribution was calculated using D’Agostino and Pearson, Anderson-Darling, Shapiro-Wilk, and Kolmogorov-Smirnov tests. If one test determined non-normally distributed data in any cell type, then non-parametric testing was used. With regard to statistical analyses, either a Mann-Whitney test with Tukey’s post hoc test was completed when comparing two means, or a Kruskal-Wallis test with Dunn’s multiple comparisons was completed when evaluating three or more means.

**Equation 2**. Slovin’s Formula was used to determine an appropriate sample size for data analysis.^92^ The total population was defined as the total number of cell bodies analyzed in each condition, and a standard error tolerance of 0.05 was used to complete the calculation. A standard value of 223 cell bodies was determined to be a significant number of samples and, therefore, was used across all conditions for continuity purposes.

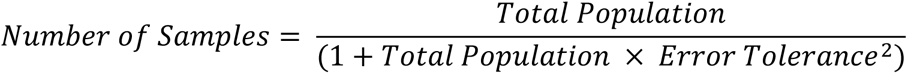

## Abbreviations

A: Area
C: Compactness
Ca^2+^ Mg^2+^: Calcium and Magnesium
CCCP: Carbonyl cyanide m-chlorophenyl hydrazone
CO_2_: Carbon Dioxide
Cy5: Cyanine-5
Drp1: Dynamin Related Protein 1
E: Eccentricity
ETC: Electron Transport Chain
FBS: Fetal Bovine Serum
glycoATP: Glycolytic ATP Production
HD: Huntington Disease
mitoATP: Mitochondrial ATP Production
MMAS: Mitochondrial Morphology Assessment System
MTR: MitoTracker ^TM^ Red
NF: Normalization Factor
NT: No Treatment
OXPHOS: Oxidative Phosphorylation
PKMDR: PKmito^TM^ Dye Deep Red
PKMR: PKmito^TM^ Dye Red
*s^2^*: variance
TIFF: Tagged Image File Format
TMRE: Tetramethyl-rhodamine Ethyl Ester Perchlorate
TruHD: Transformed HD patient fibroblasts

## Author Contributions

KD carried out the experiments and developed the pipelines presented in this manuscript. KD and RT designed the project experiments, and TM provided technical assistance. KD generated the figures and wrote the manuscript. TM, SLD, CP, and RT edited the writing while RT obtained funding.

## Data Availability Statement

All original data files and CellProfiler^TM^ and KNIME^®^ pipelines are available on Borealis using https://doi.org/10.5683/SP3/VII0UC under the Creative Commons license CC BY 4.0.^26^

## Acknowledgements

We sincerely thank Dr. Daniel J Rees (Swansea University, United Kingdom) for his help with the initial learning curve associated with CellProfiler^TM^ and Dr. Zhixing Chen (Peking University, China) for generously providing the PKmito^TM^ Dyes used for throughout this study.

